# Unexpected termite inquilines in nests of *Constrictotermes cyphergaster* (Silvestri, 1901) (Blattodea: Isoptera)

**DOI:** 10.1101/600528

**Authors:** Diogo Costa, Alessandra Marins, Og DeSouza

## Abstract

The heterospecific termite-termite cohabitation in a single termitarium, so called inquilinism, is a common event whose basal mechanisms remain hypothetical. While some termite hosts have plenty of inquilines, others house only a few. Among these, *Constrictotermes cyphergaster* are frequently found cohabiting with a single obligate inquiline species but have been unknown to house any facultative inquilines. Here we present the first record of facultative inquilines (*Embiratermes festivellus, Nasutitermes kemneri*, *Obitusitermes bacchanalis* and *Subulitermes*) to this host, evidencing that this was unlikely to have happened fortuitously. In an attempt to pose hypotheses on the mechanisms behind such invasions, we explored likely connections between the settlement of obligate and facultative inquilines and nest wall’s physical traits. We found that nests bearing atypical external walls (moist, eroded, and partially covered by mosses) held higher richness of facultative inquilines than nests presenting walls void of such traits (*χ*^2^ = 8.3965, 1 df, n = 17, P = 0.0038). The presence of healthy host colonies in all nests, including the atypical ones, reinforces the hypothesis that the settlement of these facultative inquilines depends less on host colonies biotic status and more on abiotic features associated to the nest. In addition, the presence of obligate inquilines was not affected by the nest wall status (*χ*^2^ = 8.3965, 1 df, n = 17, P = 0.0038), implying that invasion by facultative and obligate inquilines in these nests would obey distinct restrictions. While warning that these hypotheses require further testing, we suggest that their understanding could shed light on the determinants of cohabitation not only in *C. cyphergaster* but in termites in general. cohabitation interspecific interaction nest invasion barriers

## 1 Introduction

The invasion of a termite nest by another termite species, so called *inquilinism*, is a common and intriguing phenomenon, whose underlying mechanisms we are only starting to understand. While some termites species, such as *Cornitermes cumulans* (Termitidae: Syntermitinae), can coexist with many inquiline species simultaneously, others are very restrict, hosting none or only a few inquilines. Among these latter, *Constrictotermes cyphergaster* (Termitidae: Nasutitermitinae) provides an emblematic case of inquilinism. Despite extensively studied in regard to inquilinism, it was so far known to host only two inquiline species, *Inquilinitermes fur* and *Inquilinitermes microcerus* (Termitidae: Termitinae), which never occur simultaneously in the same host nest (Mathews, 1977). Other termite species frequently reported as inquilines are not found in *C. cyphergaster* nests despite overlapping biogeographical ranges. So far, this builder has thus been considered highly refractory to inquilinism.

Here we add evidence to the notion that barriers to inquilinism in *C. cyphergaster* are indeed surpassable, reporting for the first time on the presence of facultative inquilines in its nests. Based on our field observations, we also present candidate hypotheses suggesting connections between the settlement of inquilines and the current status of their host colony or their nests’ physical structure.

## 2 Material and Methods

This study was carried out in the Brazilian *Cerrado*, an environment physiognomically but not floristically similar to savannas, near the town of Diamantino (14°24’32”S, 56°26’45”W, 250 m above sea level), Mato Grosso State, Brazil, where Köppen’s Aw climate (equatorial with dry winter) prevails (Kottek et al, 2006).

*Constrictotermes cyphergaster* (Silvestri, 1901) is a common termite species in Brazil, Paraguay, Bolivia, and Northern Argentina. Its mature nests are typically arboreal, occurring more frequently on the sunnier (hence drier) side of the tree (Leite et al, 2011). Seventeen arboreal nests built by *C. cyphergaster* were sampled in March 2014 and April 2016, and notes have been taken on the current status of the external surface of these nests’ walls. Specifically, we noted any sign interpretable as a deviation from the normal dry nest aspect, such as the presence of mosses, lichens, and algae which are known to occur in termitaria under tree shade (Aptroot and Caceres, 2014).

Nests were then brought to the lab, broken into pieces, and searched to collect host and inquiline individuals with the help of a soft forceps. Sampled individuals were identified to species, following Mathews (1977) and by comparison with the collection of the Isoptera Section of the Entomological Museum of the Federal University of Viçosa (MEUV), where voucher specimens were deposited.

Pearson’s chi-square test was applied to the data, aiming to check for the independence between the atipicity of nest walls (i.e., the presence or absence of mosses) and the type of inquilines (facultative or obligatory) therein found.

## 3 Results

The 17 sampled nests were arboreal and scattered over a 9.7 ha rectangle bounded by the coordinates 14°28’ 07” S × 56°29’ 52” W and 14°28’ 22” S × 56°29’ 45” W. Nest volumes ranged from 6.14 to 34.96 L, averaging 21.28 L.

All nests presented the same general architecture, closely ressembling what is known for *C. cyphergaster* nests (Mathews, 1977; Leite et al, 2011). These features, coupled with the presence of an active and healthy colony of *C. cyphergaster*, supported attributing to this species the role of primary builder of such nests. Six of such nests were almost entirely covered by a mixture of mosses, algae and lichens. The other 11 nests presented no sign of such a coating on their external walls (Fig. 1).

**Figure 1:**
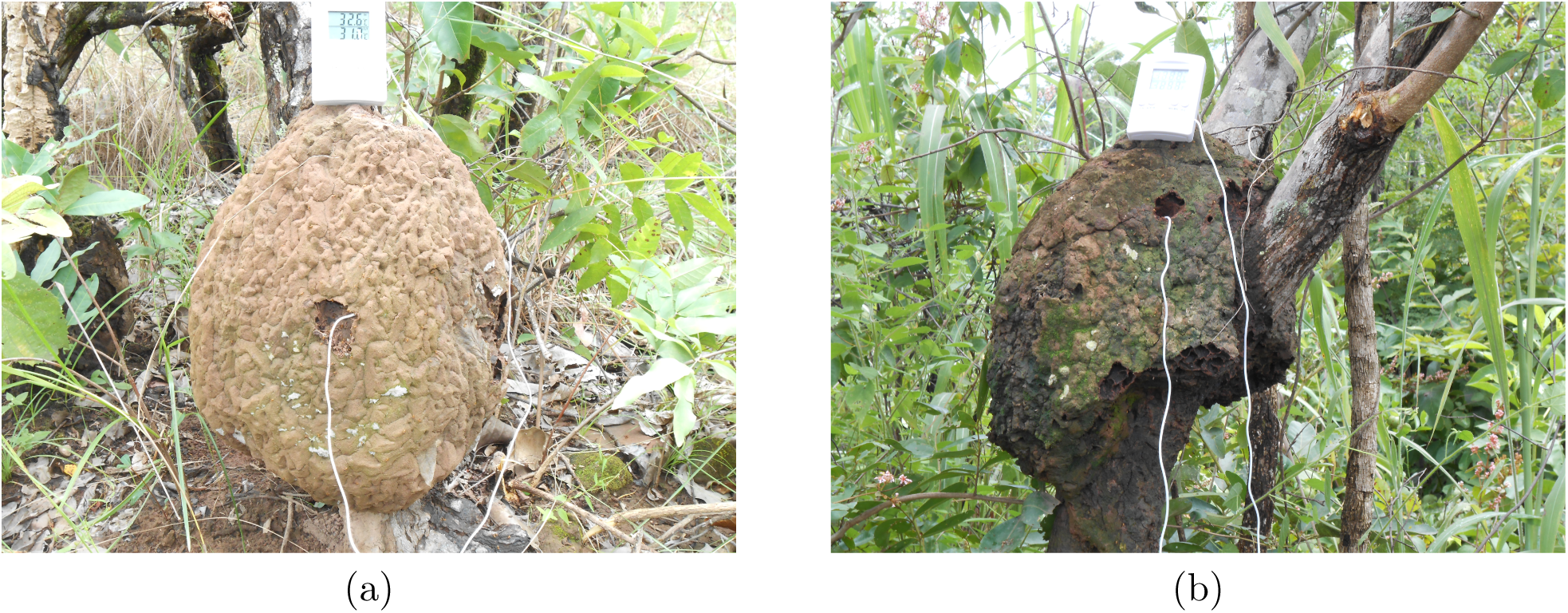
Nests of *Constrictotermes cyphergaster* sampled in Diamantino, Mato Grosso, Brazil. (a) Typical nest: light-coloured, intact, dry, and free of any coat (b) Atypical nest: dark, damaged, moist, and coated on mosses, lichens, and algae.

*C. cyphergaster* and five other termite species where found within these nests: one obligate inquiline (*Inquilinitermes fur*) and four facultative ones. The number of inquiline species in a given nest ranged from zero to three. The obligate inquiline *I. fur* was present in 13 out of the 17 nests sampled. Facultative inquilines have been found in eight nests out of the 17, seven of which also housed *I. fur*. As facultative inquilines we found *Embiratermes festivellus, Nasutitermes kemneri, Obitusitermes bacchanalis* and *Subulitermes* sp. The most frequent of them was *N. kemneri*, in six nests. *O. bacchanalis* was present in two nests and *E. festivellus* and *Subulitermes* sp. where recorded in one nest each. The most populous colonies were those of *N. kemneri* and *E. festivellus*, presenting numerous and active soldiers and workers. Four out of six colonies of *N. kemneri* also presented many nymphs. The other two species (*O. bacchanalis* and *Subulitermes* sp.), despite less populous, did not appear discrepant from what is reported as their normal (small) colony size. The facultative inquiline species here reported have been already recorded as inquilines in other termite hosts (Domingos, 1983; Redford, 1984; Costa et al, 2009; Cunha and Morais, 2010). We also found these four species in nests built by *Cornitermes bequaerti* in this same locality. A summary of these findings is given in Table 1.

**Table 1:** Inquiline termite species recorded in nests of *Constrictotermes cyphergaster* (Blattodea: Isoptera) sampled in Diamantino, Mato Grosso, Brazil, in March 2014 and April 2016. Each line refers to a single host nest, with respective field code (Nest ID), record of the presence of inquiline species on each type of nest (see Fig. 1), and nest location (latitude × longitude). All host and inquiline colonies inhabiting these nests were active and healthy. *Inquilinitermes fur* are obligate inquilines, all the others are facultative.

| Nest ID | *Inquilinitermes fur* | *Nasutitermes kemneri* | *Obtusitermes bacchanalis* | *Subulitermes* sp. | *Embiratermes festivellus* | Location |
| --- | --- | --- | --- | --- | --- | --- |
| a) Nest wall with mosses, algae or lichens |  |  |  |  |  |  |
| DM02 |  | x | x |  |  | 14°28'22" S<br>56°29'50" W |
| DM11 | x | x |  | x |  | 14°28'22" S<br>56°29'46" W |
| DM14 | x | x |  |  |  | 14°28'21" S<br>56°29'50" W |
| DM15 | x | x |  |  |  | 14°28'19" S<br>56°29'47" W |
| DM31 |  | x |  |  |  | 14°28'07" S<br>56°29'41" W |
| DM34 | x | x |  |  |  | 14°28'15" S<br>56°29'51" W |
| b) Nest wall without mosses, algae or lichens |  |  |  |  |  |  |
| DM03 | x |  | x |  |  | 14°28'21" S<br>56°29'49" W |
| DM06 | x |  |  |  |  | 14°28'22" S<br>56°29'49" W |
| DM08 | x |  |  |  |  | 14°28'19" S<br>56°29'45" W |
| DM10 | x |  |  |  | x | 14°28'21" S<br>56°29'52" W |
| DM12 |  |  |  |  |  | 14°28'19" S<br>56°29'46" W |
| DM13 | x |  |  |  |  | 14°28'21" S<br>56°29'46" W |
| DM30 | x |  |  |  |  | 14°28'07" S<br>56°29'41" W |
| DM32 | x |  |  |  |  | 14°28'10" S<br>56°29'44" W |
| DM33 | x |  |  |  |  | 14°28'14" S<br>56°29'50" W |
| DM36 |  |  |  |  |  | 14°28'13" S<br>56°29'44" W |

The most obvious feature of six out of the eight nests housing the unsual inquilines (that is, non-*Inquilinitermes*) here reported was a coat of mosses, algae, and lichens over their external walls. This atypical nest walls were darker and moister than the walls typically composing the nests of this builder. The remaining 11 nests presented no signs of mosses on their external walls (Fig. 1) and where void of these unsual inquilines. Pearson’s chi-square test with Yates’ continuity correction revealed that the existence of this coat on the nest walls was correlated to the presence of the facultative inquilines (*χ*^2^ = 7.4062, df = 1, P = 0.007) but not to the presence of the obligatory inquiline *I. fur* (*χ*^2^ = 0.0112, df = 1, P = 0.916).

## 4 Discussion

It seems likely that the inquiline records here reported were not fortuitous, at least for *N. kemneri*, which were found forming active and populous colonies in six nests of *C. cyphergaster* (Table 1), four of these colonies containing nymphs. The same might be said even for the other three species which, despite occurring only twice (*O. bacchanalis*) or once (*E. festivellus* and *Subulitermes* sp.), presented very active colonies within their host nest. Moreover, since all four invading species are typically soil-dwellers (Mathews, 1977) and given that all host nests were arboreous, their occurrence as active colonies in such nests seems not entirely accidental.

Finding such species as inquilines is not at all surprising since all them have been already recorded in termitaria in this region and elsewhere (Domingos, 1983; Redford, 1984; Costa et al, 2009; Cunha and Morais, 2010). What is surprising is their record in nests built by *C. cyphergaster*, as this opposes previous assumptions on restrictions of this host to inquilinism. Such assumptions derive, correctly, from the paucity of inquiline species not only in *C. cyphergaster* but also in other Nasutitermitinae.

The exceptionality of these findings sustains the notion that *C. cyphergaster* presents marked refractoriness to inquilines but, at the same time, it denounces that this is not an immutable trait. The reasons explaining these findings remain, however, hypothetical. After all, this same exceptionality implies that any dataset on this issue would hardly suffice for proper test of hypotheses. In spite of that, our data provide instigating hints.

One can not ascertain whether these atypical walls were a cause, a consequence, or simply coincidental of inquiline invasions in such nests. The simplest hypothesis would pose that these wall traits indicate an unhealthy colony unable to keep up the nest and to repel invaders, but the presence of populous and active host colonies in all nests seems to weaken this hypothesis. An alternative hypothesis would consider that the invasions were eased by the walls entering decayment due to external factors (such as tree shade) not necessarily connected to the host colony status. This hypothesis finds support on recent findings that pure physical nest attributes (such as size and spatial location) can be the strongest predictors of cohabiting termite species richness and composition in *C. cumulans* (Marins et al, 2016; Monteiro et al, 2017) and important determinants of invasions by obligatory inquilines and termitophiles in *C. cyphergaster* (Cristaldo et al, 2012; DeSouza et al, 2016).

Clearly, all these hypotheses require testing, and we present them to highlight research pathways that might lead to a better understanding of the phenomenon of cohabitation in nests of *C. cyphergaster* and maybe in termite nests in general. These hypotheses appear to follow naturally from the study reported here because our focal inquilines seemed not to occupy these host nests at random. Instead, nest invasion was apparently dependent the ecological context under which these nests become suitable for cohabitation.

## 5 Acknowledgements

We thank H.V. Ribeiro for his support during the field work. All necessary permits were obtained for the described study, which complied with all relevant regulations of Brazil. ODS holds a Fellowship from National Council for Scientific and Technological Development (CNPq PQ 307990/2017-6). Computational work was performed using free software (GNU-Linux/Debian and Ubuntu, LaTeX, Gimp, Kile, LibreOffice, RStudio, and R, among others). This is contribution no. # from the Termitology Lab at UFV, Brazil (http://www.isoptera.ufv.br).

